# A timed epigenetic switch balances T and ILC lineage proportions in the thymus

**DOI:** 10.1101/2024.05.04.592518

**Authors:** Nicholas Pease, Lihua Chen, Peter Gerges, Hao Yuan Kueh

## Abstract

How stem and progenitor cells give rise to multiple cell types in defined numbers and proportions is a central question in developmental biology. Epigenetic switches, acting at single gene loci, can generate extended delays in the activation timing of lineage-specifying genes, and thereby impact lineage decisions and cell type output of progenitors. Here, we analyzed a timed epigenetic switch controlling *Bcl11b*, a transcription factor that drives T cell lineage commitment, but only after a long multi-day time delay in expression. To investigate roles for this delay in controlling lineage decision making, we analyzed progenitors with a deletion in a distal *Bcl11b* enhancer, that further extends this delay by ∼3 days. Strikingly, delaying *Bcl11b* activation reduces T cell output but enhances ILC generation in the thymus, and does so by redirecting progenitors to the ILC lineages at the T and ILC developmental branchpoint. Mechanistically, delaying *Bcl11b* activation promoted ILC redirection by up-regulating a PLZF-dependent ILC transcriptional program in progenitors. Despite up-regulating PLZF, committed ILC progenitors were still capable of later activating *Bcl11b*, which is also needed for specification of type 2 ILCs. These results show that epigenetic switches, by controlling the activation timing and order of lineage-specifying genes within regulatory networks, can modulate population sizes and proportions of differentiated cell types.

## INTRODUCTION

During development and tissue regeneration, stem and progenitor cells give rise to differentiated cell types in defined numbers and proportions to properly generate and maintain organs and body plans. To do so, these cells must closely control their lineage decisions in space and in time inside an embryo or organism. The timing of lineage decisions, in particular, is subject to close developmental control, as it shapes both the degree to which progenitors can proliferate, as well as their internal regulatory state and lineage potential.

Epigenetic switching mechanisms, which modulate the activation timing of lineage-specifying genes, can influence stem cell lineage decision making (Abadie et al., 2019; Ebisuya and Briscoe, 2018). The activation timing of genes is often assumed to depend solely on the activity of upstream *trans*-factors. However, across a range of systems, epigenetic switches, acting in *cis* at individual gene loci, can delay the activation of lineage-specifying genes after initial exposure to developmental signals (Abadie et al., 2023; Berry et al., 2017; Bintu et al., 2016; Hathaway et al., 2012; Ng et al., 2018), sometimes by multiple days and cell generations. These time delays in epigenetic switching vary both between single cells and between individual gene loci in the same cell due to the stochastic nature of chromatin regulation (Bintu et al., 2016; Dodd et al., 2007; Festenstein et al., 1996; Owen et al., 2023) However, despite this stochasticity, probabilistic time constants of switching are precisely modulated at multiple levels, including by cytokine signaling, transcription factors (TFs), and associated *cis*-regulatory elements and chromatin-modifying enzymes (Chu et al., 2021; Deschamps and Duboule, 2017; Fabre et al., 2015; Kissiov et al., 2022; Pease et al., 2021). Because of their tunable nature, such timed epigenetic switches could be utilized by stem cells to control differentiation output. However, it has remained unclear what roles, if any, timing delays in epigenetic switching play in controlling stem cell lineage decisions and output.

Here, we study the functional roles for epigenetic timing delays in the lineage decisions of early thymic progenitors. After entering the thymus, hematopoietic progenitors maintain a multipotent state for ∼7 days, where they are able to give rise to multiple immune cell lineages. The large majority of progenitors then commit to the T cell lineage, and proceed to undergo T cell receptor rearrangement and selection to become a functional T cell. However, there is a fraction of progenitors that can give rise to type 2 innate lymphoid cells (ILC2s) as an alternate lineage option, particularly in the embryonic or neonatal thymus^18^, or generate distinct subtypes of natural killer (NK) cells^21,22^.

The extended time period in which progenitors maintain a multipotent state is due in part to an epigenetic switch that generates a multi-day delay in the activation of the T cell lineage commitment regulator *Bcl11b*. T cell lineage specification begins with engagement of Notch/Delta signaling (Chen et al., 2019; Maillard et al., 2005; Radtke et al., 2010), which up-regulates the expression of the transcription factors TCF-1 and Gata3. These TFs, in conjunction with Runx factors that are already expressed prior to thymic entry, then work together to turn on Bcl11b, which represses myeloid and NK cell potential and drives T cell commitment (García-Ojeda et al., 2013; Germar et al., 2011; Kueh et al., 2016; Shin et al., 2023; Weber et al., 2011). Strikingly, while TCF-1 and Gata3 turn on shortly after thymic entry and exposure to Notch ligands (∼1 day), Bcl11b activation occurs only ∼7 days later (Rothenberg, 2019; Yui and Rothenberg, 2014). By analyzing *Bcl11b* activation dynamics at the single chromosomal allele / single cell level, we found that this multi-day delay was due to an timed epigenetic switch, involving a rate-limiting transition of the gene locus from a compacted, silent state to a de-compacted, expressing state (Ng et al., 2018; Pease et al., 2021). Importantly, while this switch is stochastic, time constants for probabilistic switching are tightly controlled by the histone-modifying enzymes [Polycomb Repressive Complex (PRC)2] (Pease et al., 2021), upstream transcription factors (TCF-1, Gata3)(Kueh et al., 2016), as well as a far-distal *cis*-regulatory element (Li et al., 2013; Ng et al., 2018), which we termed a ‘timing enhancer’ as it moderately speeds up the onset of *Bcl11b* activation in progenitors but is not required for maintaining its expression (Chu et al., 2021).

Time-delayed control of *Bcl11b* epigenetic switching and activation may play a role in T cell and ILC2 lineage decision making in early progenitors. While ILCs share common transcriptional programs with T cells, they also express Id2, PLZF (encoded by *Zbtb16*) and RORα, which work together to suppress T cell programs (Qian et al., 2019; Wang et al., 2017) and to specify ILC identity (Constantinides et al., 2014; Fang and Zhu, 2017; Ferreira et al., 2021; Rothenberg, 2019). These ILC-specific regulators are prevented from being expressed in committed T cell progenitors by expression of Bcl11b (Hosokawa et al., 2018; Zhou et al., 2022); however, because these regulators are downstream of Notch signaling and its target Gata3 (Ferreira et al., 2021; Rothenberg, 2019), they could potentially become induced in multipotent progenitors, particularly in cells that show lengthened delays in *Bcl11b* activation. Paradoxically, although Bcl11b represses Id2 and PLZF, its expression is also required for development of ILC2s, where it binds and regulates a set of genes distinct from its target genes in T cells (Hosokawa et al., 2020). It is mysterious as to how the same TF that represses alternate-fate regulators to uphold cell identity in one lineage, can also work alongside the same regulators to establish another identity in another lineage.

In this study, we seek to understand how developmental timing delays set by the *cis*- epigenetic switch for *Bcl11b* activation regulate T and ILC decisions in early DN progenitors. To disentangle *cis*-epigenetic timing control from other developmental mechanisms, we utilized a knockout mouse strain carrying a deletion of the *Bcl11b* timing enhancer (Li et al., 2013; Ng et al., 2018) (ΔTE). TE serves as one of two transcription start sites for a distal long non-coding RNA transcript (*ThymoD*) which is activated immediately preceding *Bcl11b* activation and physically interacts with its promoter through DNA looping mechanisms (Isoda et al., 2017). While completely disrupting *ThymoD* transcription impairs *Bcl11b* expression levels and leads to impaired T cell development and consequently leukemia, removal of TE only moderately delays *Bcl11b* activation by ∼3 days without affecting its maintenance and subsequent T cell maturation and function once expressed. By analyzing lineage decision making in cells having both normal and protracted time delays in *Bcl11b* activation in the thymus and *in vitro* (Holmes and Zuniga-Pflucker, 2009), we found delayed *Bcl11b* activation and T cell lineage commitment reduced T cell output, but enhanced the generation of ILC2s in the thymus. This lengthened time in an uncommitted state enabled progenitors to prime an early pro-ILC transcriptional program, marked by heightened expression of PLZF, which drove commitment down the ILC2 pathway. Importantly, progenitors were still able to activate *Bcl11b* after the onset of ILC priming; however, when *Bcl11b* was activated after priming, it no longer repressed PLZF and other ILC2 regulators, which were now stably co-expressed alongside *Bcl11b* to enable the commitment to the ILC2 fate. These results show that *cis*-epigenetic switches, by setting the relative timing of gene regulatory events, can control lineage decisions at a developmental branch point and thereby control the numbers and proportions of differentiated cells generated. More generally, our findings highlight the importance of temporal regulation in gene regulatory networks for decision making in development and tissue regeneration.

## RESULTS

### A distal enhancer modulates the activation timing of *Bcl11b* but not maintenance of its expression

It is generally difficult to perturb the timing of a single epigenetic switching event and lineage commitment step in isolation from other molecular processes in the cell. Knocking out an essential lineage-specifying gene would generally completely abrogate development, whereas perturbing upstream signals or *trans*-acting factors that regulate the gene of interest would lead to systemic regulatory effects that are difficult to disentangle from effects on target gene activation itself.

Mutating a *cis*-regulatory element that selectively modulates the kinetics of a gene activation event *in cis* provides a method for probing functional roles for timing in lineage decision making. From work in diverse developmental and differentiation systems, it is now established that some *cis*-acting regulatory elements specifically enhance the probability a single gene stochastically switches on in response to signals, without affecting its final expression magnitude once active. As switching probabilities are low, they can frequently give rise to extended gene activation delays that span multiple cell generations, and thus can be referred to as “timing enhancers” (Chu et al., 2021). Mutations at these timing enhancers could provide a unique tool to investigate roles for gene activation timing (Nguyen et al., 2021), as they frequently result in moderate changes in the timing delays for gene activation and developmental events (Gérard et al., 1997; Zákány et al., 1997).

To study the role of *Bcl11b* activation timing in regulating thymocyte differentiation, we utilized a mouse in which the *Bcl11b* timing enhancer was removed from both copies of *Bcl11b*, which were also non-disruptively tagged with a mCitrine yellow fluorescent protein (Bcl11b^YFP-ΔTE^) (Ng et al., 2018). To first assess the effects of timing enhancer deletion on *Bcl11b* activation dynamics, we crossed this Bcl11b^YFP-ΔTE^ mouse strain to a second strain in which each wild-type copy of *Bcl11b* was non-disruptively tagged with a mCherry red fluorescent protein (Bcl11b^RFP-WT^). We purified bone marrow derived Bcl11b-negative DN2a progenitors from either Bcl11b^YFP-WT/RFP-WT^ or Bcl11b^YFP-ΔTE/RFP-WT^ mice and co-cultured them with OP9-DL1 stromal cells, which enable the reconstitution of early thymopoiesis *in vitro*. In agreement with our previous results(Ng et al., 2018), the mutant Bcl11b^YFP-ΔTE^ allele was expressed in an all-or-none manner ∼3 days later than the wild-type Bcl11b^RFP-WT^ allele in the same cells (Figure 1A, Figure S1A). Similarly, Bcl11b^YFP-ΔTE^ DN2a progenitors exhibited a delayed onset of YFP expression relative to Bcl11b^YFP-WT^ DN2a progenitors; however, Bcl11b^YFP-ΔTE^ DN2a progenitors were able to maintain relatively normal Bcl11b-YFP expression levels after activation (Figure 1A). To determine whether this timing enhancer mutation could be used to study the importance *Bcl11b* activation timing *in vivo*, we next compared Bcl11b expression across all stages of thymocyte development in Bcl11b^YFP-WT/YFP-WT^ (WT) or Bcl11b^YFP-ΔTE/YFP-ΔTE^ mice (ΔTE) (Figure 1B-E). ΔTE mice exhibited a reduction in the percentage of Bcl11b-YFP+ DN2 progenitors, confirming that Bcl11b activation is delayed at the DN2 stage *in vivo* (Figure 1D). However, as observed *in vitro*, ΔTE thymocytes at the DN3-DP stages maintained comparable levels of Bcl11b expression post-activation and thymocyte maturation (Figures 1C, 1E). Together, these results demonstrate that removal of the *Bcl11b* timing enhancer selectively delays *Bcl11b* activation at the DN2 stage without disrupting *Bcl11b* expression levels and subsequent stages of T cell development.

**Fig. 1:**
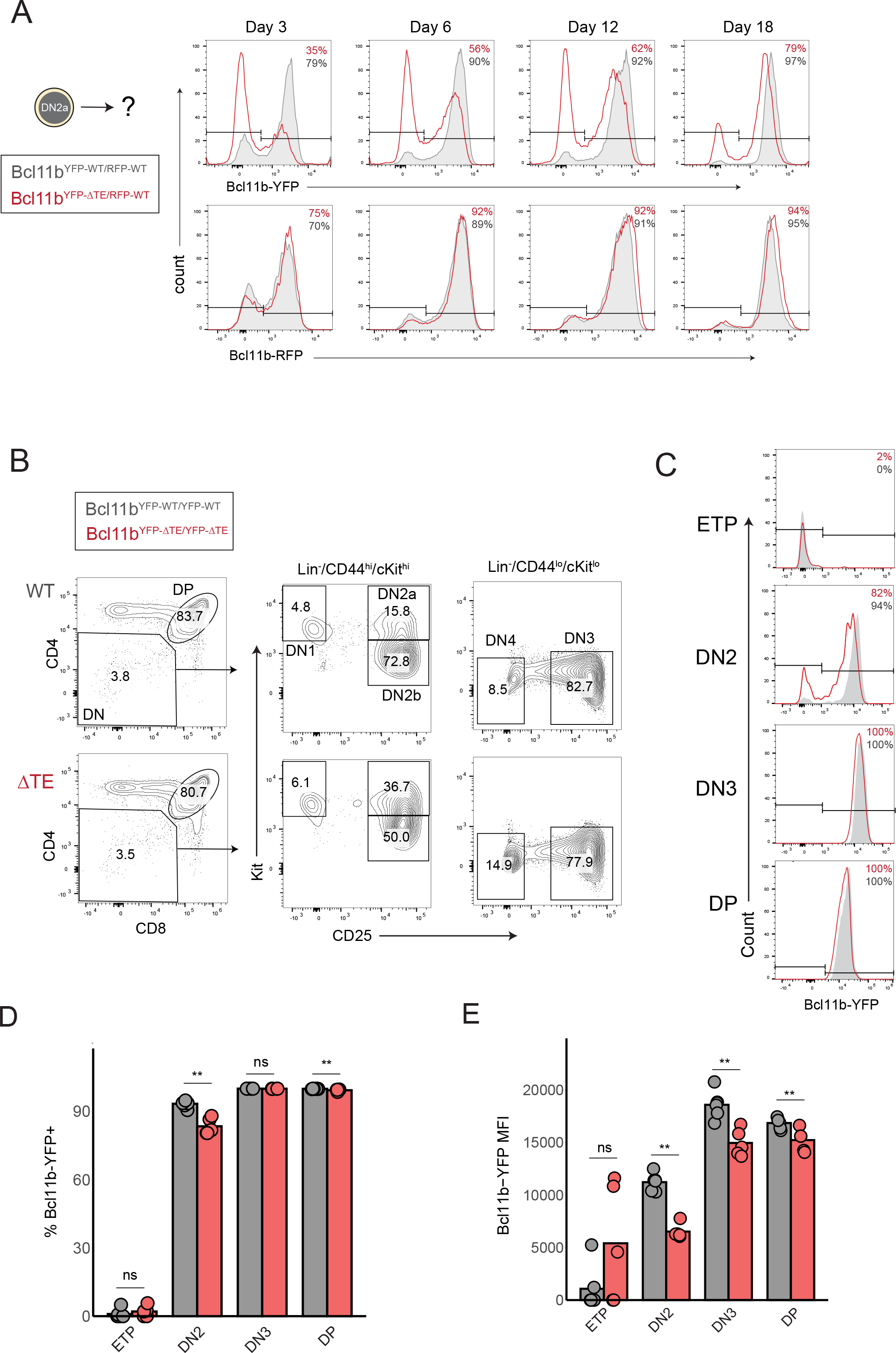
Deletion of a distal enhancer selectively delays the timing of *Bcl11b* activation without altering its expression magnitude or maintenance. (A) Bone marrow derived Bcl11b-negative DN2a progenitors were sorted and re-cultured on OP9-DL1 stromal cells for up to 18 days. (B) Gating strategy for analyzing *in vivo* thymocytes. (C) Representative histograms of Bcl11b-YFP expression levels for each T cell developmental stage. (D) Quantification of frequency of Bcl11b- YFP+ thymocytes (Wilcoxon rank sum test,, **p < 0.01, n = 5-6 mice per genotype). (E) Quantification of Bcl11b-YFP expression levels thymocytes (geometric mean fluorescence intensity; Wilcoxon rank sum test, **p < 0.01, n = 5-6 mice per genotype).

### Delayed *Bcl11b* activation decreases T cell output and increases thymic ILC output

To investigate the importance of *Bcl11b* activation timing in T cell development, we compared stage-specific T cell frequencies and numbers in thymi of wildtype and ΔTE mice. We found that delayed *Bcl11b* activation in ΔTE mice significantly increased the percentage of pre-committed DN2a thymocytes (∼1.7-fold) yet did not disrupt T cell development and maturation after lineage commitment (Figure 2A), consistent with a delay in T cell lineage commitment at the DN2 stage upon TE deletion. This extended delay in T cell lineage commitment resulted in significant decreases in the total number of double-positive (DP) thymocytes (-16%) (Figure 2B), as well as peripheral T cells in the spleen (-27%) (Figure 2C). Importantly, the number of B cells in the spleen remained constant (Figure 2C), suggesting that these changes in T cell numbers are not due to systemic changes in immune cell development in ΔTE mice. These results indicate that a delay in the timing of T cell lineage commitment, due to deletion of the *Bcl11b* timing enhancer, leads to an attenuation in T cell output from the thymus.

**Fig. 2:**
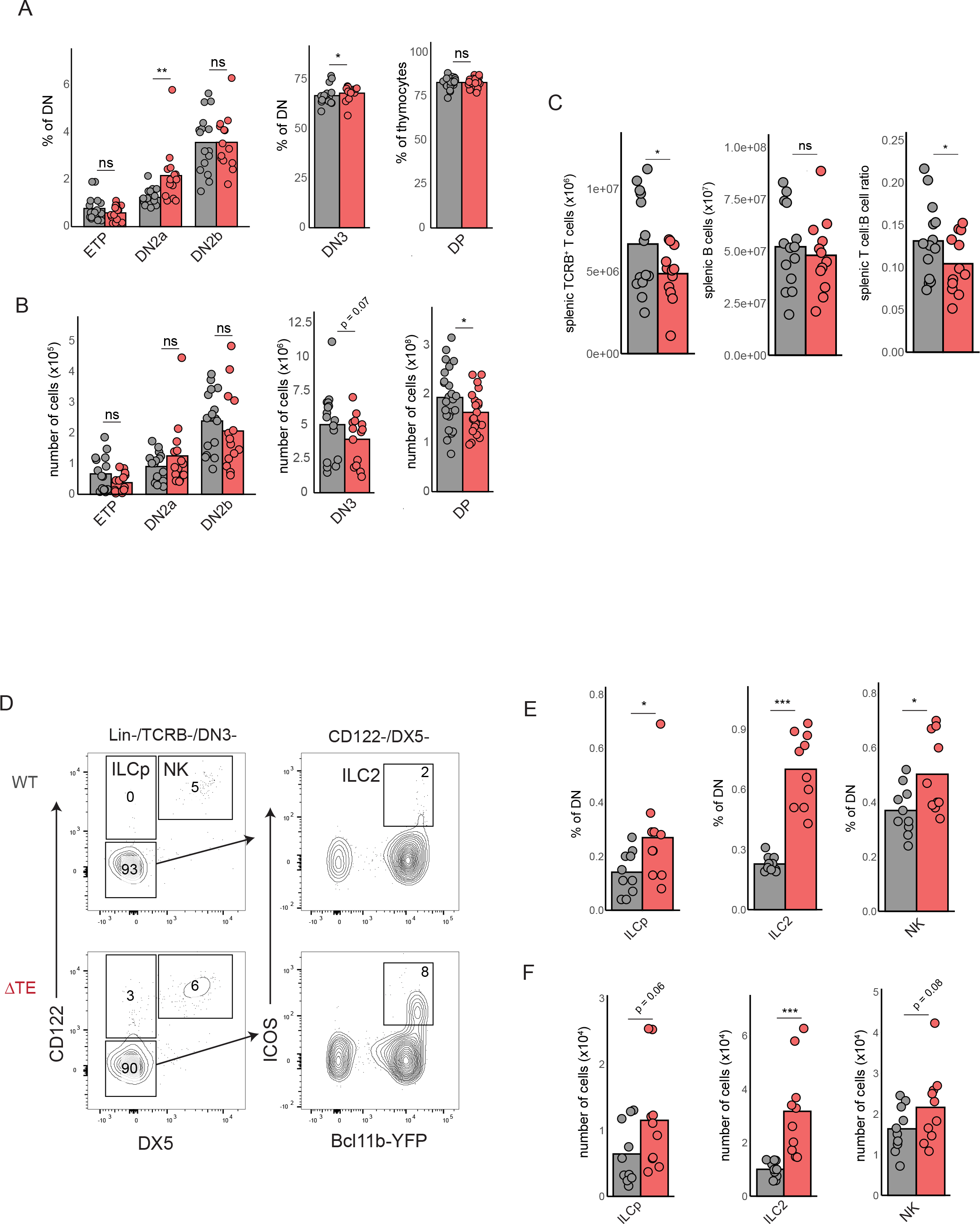
Delaying *Bcl11b* activation enhances ILC generation while reducing T cell output. (A-B) Quantification of thymocyte population frequencies and total cell numbers (Wilcoxon rank sum test, *p < 0.05, **p < 0.01, n = 15-16 per genotype for DN populations and n = 22-24 per genotype for DP cells.) (C) Quantification of splenocyte cell numbers (T-test, *p < 0.05, n = 12- 14 per genotype). (D) Gating strategy for analyzing thymic ILC subsets. (E-F) Quantification of thymic ILC subset frequencies and total cell numbers (Wilcoxon rank sum test, *p < 0.05, *** <0.001, n = 10 per genotype).

Activation of *Bcl11b* drives the silencing of stem cell and non-T cell lineage specifying genes in progenitors^22,36^ and it’s deletion results in the acquisition of an immature ILC-like state, suggesting that innate lymphoid cell lineage is the final non-T cell lineage that is restricted upon expression of Bcl11b at the DN2 stage (Hosokawa et al., 2018; Li et al., 2010; Zhou et al., 2022). Therefore, delaying *Bcl11b* activation may increase the frequency of progenitors that diverge away from the T cell lineage and towards the ILC pathway in the thymus. To investigate this hypothesis, we quantified the frequency of ILC precursors (ILCp) (Lin-/TCRB-/CD122+/DX5-), ILC2s (Lin-/TCRB-/CD122-/DX5-/ICOS+/Bcl11b+) and NK (Lin-/TCRB-/CD122+/DX5+) cells in thymi of wildtype and ΔTE mice (Figures 2D-2F). Consistent with their status as immature precursors, ILCp cells retained heterogenous levels of uncommitted progenitor markers c-Kit, PD-1, and CD44 (Figure S2A-B). Furthermore, mature NK and ILC2 cells were low for c-Kit and CD25 with heterogeneous levels of PD-1 and CD44 (Figure S2A-B), in agreement with previous studies (Rodriguez-Rodriguez et al., 2022). Compared to the wildtype mice, ΔTE mice exhibited significantly elevated frequencies and numbers of ILCp and ILC2 cells in the thymus; they also exhibited a moderate increase in the frequencies of NK cells. Together, these results indicate that a prolonged delay in the activation timing of *Bcl11b* can result in enhanced production of multiple ILC lineages. Notably, ILC2s generated from ΔTE progenitors expressed Bcl11b at similar levels compared to those generated from wildtype progenitors. This similarity indicates that differences in ILC2 numbers between wildtype and ΔTE mice arise due to differences in activation timing and not dosage differences in *Bcl11b* expression. Together, these results demonstrate that the timing of Bcl11b activation is important for determining the relative proportions of T cell and ILC progenitors in the thymus.

### Delayed *Bcl11b* activation promotes ILC diversion in multipotent thymic progenitors

Previous studies had shown that thymic progenitors retain the capacity to differentiate into ILCs for multiple days after thymic entry, prior to *Bcl11b* activation and T cell lineage commitment (Qian et al., 2019; Wang et al., 2017). These observations raise the possibility that the delay in *Bcl11b* activation due to timing enhancer deletion may promote ILC diversion at multiple stages leading up to T cell commitment by prolonging the time uncommitted progenitors spend in this multipotent state. To directly test this hypothesis, we compared the ILC lineage potential of wildtype versus ΔTE progenitors at multiple stages, both before and after T cell lineage commitment. To do so, we sorted pre-commitment (ETP and DN2a) and post-commitment (DN2b and DN3) progenitors from the thymi of wildtype and ΔTE mice, co-cultured them with OP9-DL1 stromal cells and cytokines to facilitate both ILC and T cell development for seven days before analyzing them byflow cytometry (Figure 3A). As expected, ETPand DN2a progenitors from wildtype mice gave rise to significant populations of ILC2s and NK cells, whereas DN2b and DN3 thymocytes showed significantly decreased production of both ILC lineages, consistent with T cell lineage commitment occurring at the DN2b stage. Timing enhancer deletion decreased T cell production while increasing NK and ILC2 generation from both ETP and DN2a progenitors (Figures 3B, 3C). ILC production was most enhanced when starting from DN2a progenitors, consistent with this lineage bifurcation occurring at this later stage where cells are poised to turn on *Bcl11b*. In contrast, DN3 progenitors from ΔTE mice generated normal frequencies of T cell progenitors and failed to generate a significant number of ILC2 or NK cells, consistent with these progenitors being T cell lineage-restricted. Together, these results demonstrate that delaying the onset of *Bcl11b* activation increases the likelihood of DN2 progenitor divergence away from the T cell lineage towards the ILC pathway in a thymocyte cell-intrinsic manner.

**Fig. 3:**
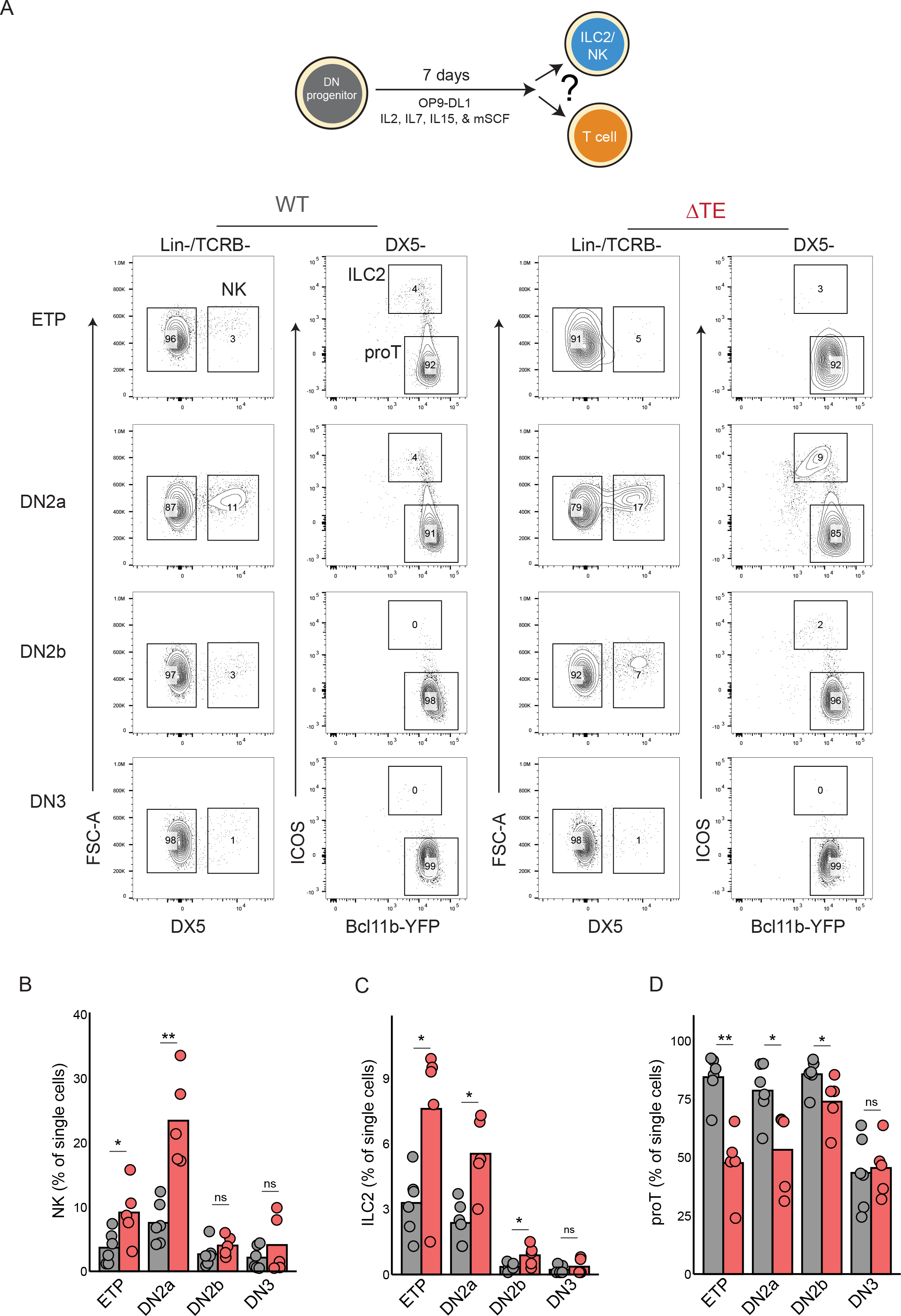
Delayed *Bcl11b* activation promotes ILC redirection in progenitors poised for T cell commitment. (A) DN thymocytes were sorted and re-cultured for 7 days on OP9-DL1 stromal cells in the presence of pro-T and pro-ILC cytokines (IL-2, IL-7, IL-15, Flt3-L, mSCF). (B-D) Quantification of the frequencies of NK, ILC2 and T cell progenitors at Day 7 (Wilcoxon rank sum test, * p < 0.05, ** p < 0.01; n = 5-6 mice per genotype) .

### Delayed *Bcl11b* activation leads to ILC priming and PLZF up-regulation in multipotent progenitors

The prolonged delay in *Bcl11b* activation due to timing enhancer deletion may promote ILC divergence by enabling a parallel and competing pro-ILC transcriptional program to emerge in DN2a progenitors. To test this hypothesis, we purified bone marrow-derived DN1 (CD44^+^CD25^-^), DN2a (CD44^+^CD25^+^Bcl11b^-^), and DN2b (CD44^+^CD25^+^Bcl11b^+^) progenitors from WT or ΔTE mice and re-cultured them on OP9-DL1 stromal cells for four days before harvesting them for single-cell RNA-sequencing (Figure 4A-C). We chose this range of initial progenitors to ensure adequate sampling of cells across different developmental states, and also to enable concurrent analysis of lineage potential for different progenitor states through sequencing. We utilized a combinatorial indexing strategy for single-cell profiling (Cao et al., 2019), which yielded high quality transcriptomes from 68,464 cells across these 6 conditions (Figure 4A-C). To clearly resolve developmental lineages from single cell data, we used a probabilistic topic modeling approach, which represents cells according to collections of co-expressed genes (i.e. gene topics) (Lynch et al., 2022). Here, each cell is represented by a mixture of gene topics, each with unique weights according to its gene expression profile (Figure S3A, Table S1). We then visualized cells based on their unique topic weights in two-dimensions using a uniform manifold approximation projection (UMAP), and clustered cells using the Leiden algorithm to visualize developmental states and trajectories.

**Fig. 4:**
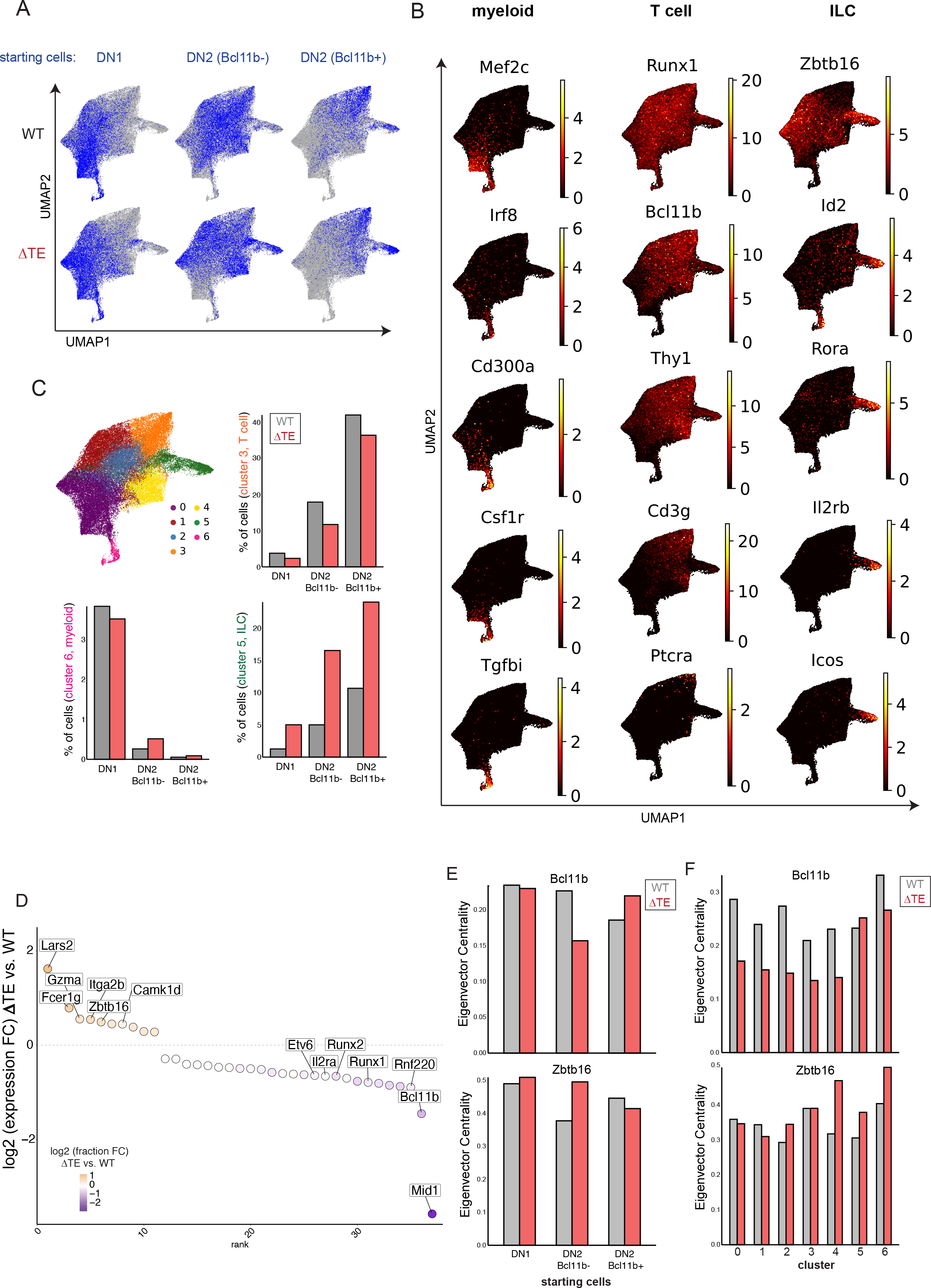
Delayed *Bcl11b* activation enables emergence of an ILC program in poised progenitors marked by *Zbtb16* up-regulation. (A) UMAP representation of scRNA-seq experiment colored by starting cell population for each sample. (B) UMAP representation of all cells colored by myeloid, T cell or ILC lineage marker gene expression levels (normalized). (C) UMAP representation of all cells colored by Leiden cluster label and quantification of myeloid, T cell and ILC cluster frequencies across samples. (D) Differential gene expression analysis between ΔTE and WT cells in the progenitor clusters (clusters 0, 1, 2, and 4 only). Only genes detected in > 5% of cells with an adjusted p-value < 0.05 and an absolute fold-change > 1.2 are displayed. (E) Eigenvector centrality values for Bcl11b and Zbtb16 in CellOracle-generated gene regulatory network models based on starting cell state or Leiden cluster label.

This visualization revealed distinct trajectories corresponding to the progression of multipotent progenitors into the myeloid, T cell or ILC lineages based on marker gene expression (Figure 4B-C). Multipotent progenitors occupied the left region of the UMAP (clusters 0, 1, 2, 4) and gave rise to three lineages, myeloid progenitors (cluster 6), DN3 T cell precursors (cluster 3) as well as ILCs (cluster 5). As expected, clusters from these lineages were enriched for expression of corresponding lineage-specific genes (e.g. *Cd3g* and *Ptcra* for T cell precursors(cluster 3); *Il2rb*, *Icos, Rora* for ILC precursors (cluster 5); *Csf1r, Mef2c* for myeloid progenitors (cluster 6). Myeloid lineages appeared to be specified early, branching off from early progenitors (cluster 0) that can also give rise to T and ILC potent progenitors; in contrast, ILC lineages appeared to be specified later, branching off progenitors (clusters 1, 2, and 4) that already appear to have some expression of T cell-lineage associated genes (*Thy1*, *Bcl11b*). These results are consistent with the ILC lineage decision point occurring late, immediately prior to T cell lineage commitment in DN2 progenitors.

Analysis of progenitor populations from WT and ΔTE mice showed enhanced ILC specification as a result of delayed *Bcl11b* activation, consistent with the results obtained from thymus-derived progenitors (Figure 3). Starting DN1 progenitors from WT and ΔTE mice generated some myeloid progenitors (cluster 6), but primarily gave rise to T cell and ILC progenitors (clusters 1, 2 and 4), reflecting transition to a T/ILC bipotent state yet to be perturbed by a forthcoming delay in *Bcl11b* activation (Figure 4C). In contrast, starting DN2 *Bcl11b*^-^ progenitors generated very few myeloid progenitors, but instead gave rise to T cell or ILC precursors, reflecting their more advanced developmental state. DN2 *Bcl11b*^-^ progenitors from ΔTE mice showed reduced T cell output compared to their wildtype counterparts (8% vs. 12%, cluster 3) but instead showed increased ILC specification (17% vs. 5%, cluster 5), consistent with the T/ILC commitment decision from these later starting progenitors being modulated with the timing of *Bcl11b* activation. Interestingly, while starting DN2 *Bcl11b*^+^ progenitors mainly gave rise to T cell precursors, as expected, they still gave rise to the significant fractions of ILC precursors, particularly for those harboring the enhancer deletion, where ILC specification was significantly enhanced (24% vs 10%). This generation of ILCs from DN2 *Bcl11b^+^* progenitors here compared to those from thymic progenitors may reflect enhanced ILC specification under *in vitro* differentiation conditions compared to the thymus. Together, these results demonstrate that DN2 progenitors retain ILC developmental potential and that delayed *Bcl11b* activation and T cell lineage commitment promotes divergence towards a pro-ILC transcriptional program within which late-coming Bcl11b expression can operate to promote ILC2 differentiation.

To identify the transcriptional regulators that promote ILC divergence in DN2 progenitors when *Bcl11b* activation is delayed, we analyzed genes differentially expressed between WT and ΔTE progenitors (clusters 0, 1, 2 and 4) that still harbor T and ILC lineage potential. From this analysis, we found that progenitors with a deleted enhancer showed significantly decreased expression of *Bcl11b*, as expected. ΔTE progenitors also showed reduced expression of other T cell lineage regulators, including *Runx1* and *Runx2*, suggesting the emergence of alternate regulatory programs that begin to dampen T cell lineage gene expression when *Bcl11b* activation is delayed (Figure 4C). Indeed, progenitors from ΔTE mice showed heightened expression of several genes associated with ILCs or NK cells, including *Fcer1g*, *Gzma* and *Zbtb16*, which encodes the pro-ILC transcription factor PLZF. Intriguingly, though ILC specification requires the concerted action of three regulators – *Zbtb16*, *Id2* and *Rora* (Constantinides et al., 2014; Fang and Zhu, 2017; Ferreira et al., 2021; Rothenberg, 2019) – only *Zbtb16* was noticeably up-regulated in uncommitted progenitors upon *Bcl11b* enhancer deletion. In contrast, *Id2* and *Rora* both showed similarly low expression levels in multipotent progenitors with a deleted *Bcl11b* enhancer, and increased in expression only after ILC pathway entry (Figure 4B).

These findings, along with observations that the *Zbtb16* locus is directly bound and repressed by Bcl11b during T cell commitment (Hosokawa et al., 2018; Hosokawa et al., 2020), implicate PLZF as a driver of ILC lineage divergence at the DN2 stage when *Bcl11b* expression is delayed. To investigate whether PLZF has elevated regulatory activity when Bcl11b is delayed, we used CellOracle gene regulatory network analysis to infer the centrality of Zbtb16 to the transcriptional programs of ΔTE and WT progenitors. CellOracle applies a linear machine-learning model to predict cluster-specific TF-to-gene linkages based on TF motif occurrence at regulatory regions associated with a target gene as well as the relationship between TF expression and that of the potential target gene (Kamimoto et al., 2023). As expected, the CellOracle model predicted Bcl11b has reduced centrality in the network of ΔTE-derived progenitors starting from the DN2 Bcl11b- state and across all progenitor clusters (0, 1, 2, 4) (Figure 4E-F). Conversely, Zbtb16 had higher centrality scores in ΔTE-derived progenitors starting in the DN2 Bcl11b^-^ state compared to their WT counterparts. Specifically, the predicted Zbtb16 centrality is most dramatically elevated in ΔTE progenitors in cells of the progenitor clusters 2 and 4 as well as the ILC cluster 5. As expected, several of the top predicted PLZF target genes were differentially expressed between ΔTE-derived and WT-derived DN2 Bcl11b^-^ progenitors (*Pde4d*, *Camk1d*, and *Runx1*) (Figure S3B). Together these findings suggest that *Zbtb16*/PLZF up-regulation in DN2 progenitors due to delayed *Bcl11b* activation may initiate a pro-ILC program in these cells.

To investigate whether delayed *Bcl11b* activation due to enhancer deletion also facilitates PLZF up-regulation *in vivo*, we measured PLZF levels in progenitors from thymi of WT and ΔTE mice using immunofluorescence staining (Figure 5A). In wild-type thymic progenitors, PLZF was low in ETPs, and was transiently upregulated at the DN2a stage before being downregulated as Bcl11b expression increased at the DN3 stage (Figure 5A), in agreement with our sci-RNA-seq results on *in vitro–*derived T cell progenitors. In progenitors from ΔTE mice, this transient expression of PLZF was elevated (∼50%) at the DN2a stage and persisted through the DN2b stage before again being down-regulated in DN3 cells. These results indicate that the ILC priming in uncommitted DN2 progenitors, as observed in *in vitro*-generated T cell progenitors, also occurs during T cell development in the thymus.

**Fig. 5:**
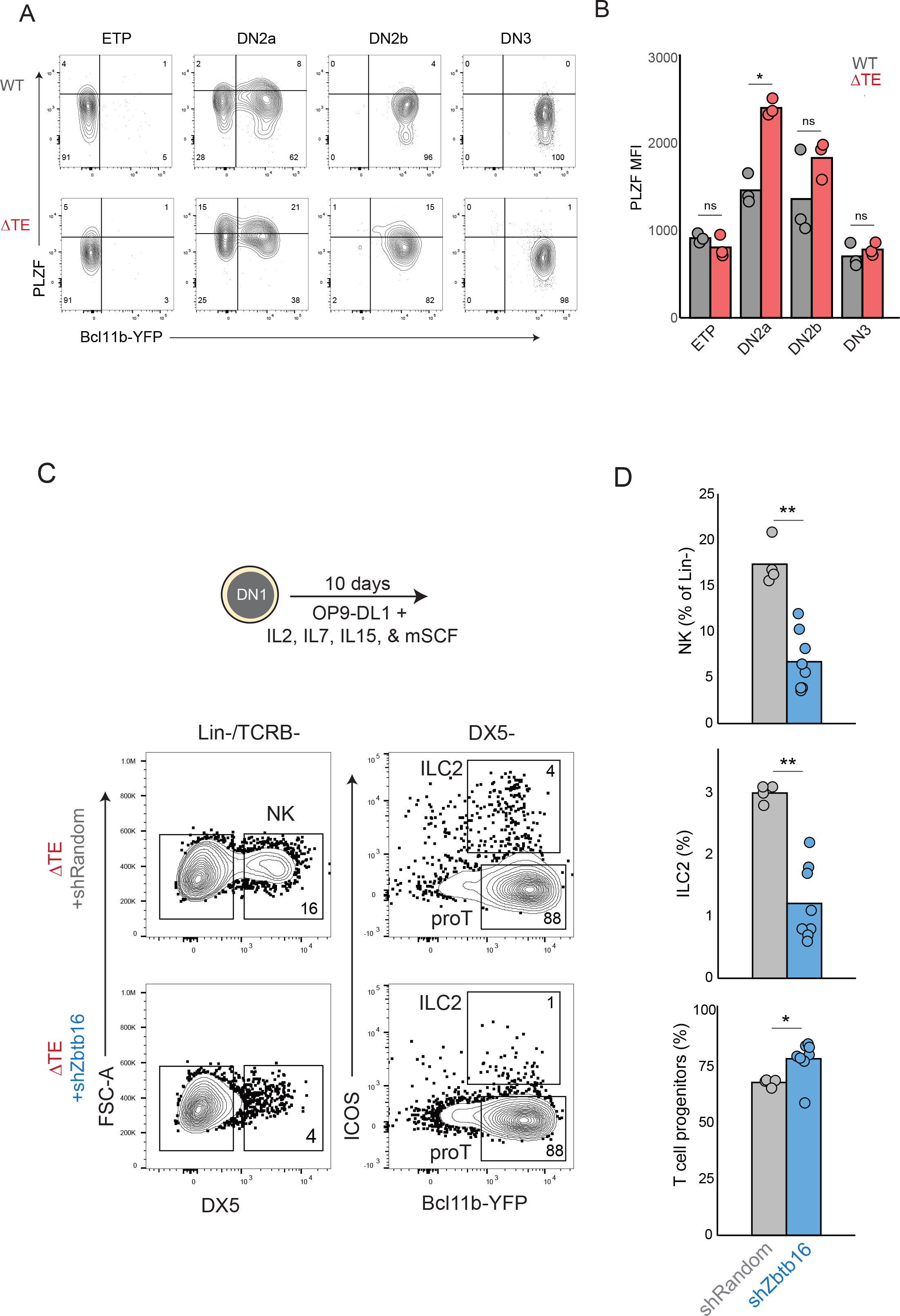
PLZF mediates ILC redirection in progenitors with delayed Bcl11b activation. (A) Representative scatter plots of PLZF and Bcl11b-YFP levels across *in vivo* DN thymocyte populations. (B) Quantification of PLZF levels (geometric mean fluorescence intensity) in thymocytes. (C) Bone marrow-derived DN1 progenitors from ΔTE mice were retrovirally transduced, sorted and re-cultured on OP9-DL1 cells in the presence of pro-ILC and pro-T cell cytokines for 10 days. (D) Quantification of the frequencies of NK, ILC2 and T cell progenitors at Day 10 (Wilcoxon rank sum test, * p < 0.05, ** p < 0.01; n = 4 (WT) or 8 (TE) mice).

### PLZF promotes ILC diversion in T cell progenitors with delayed Bcl11b activation

The heightened up-regulation of PLZF in DN2 progenitors experiencing delayed Bcl11b expression may be responsible for the increased frequency of thymic progenitor divergence into the ILC pathway. To test this hypothesis, we knocked down PLZF expression in ΔTE-derived DN1 progenitors through retroviral transduction of small hairpin (sh)RNAs targeting *Zbtb16* (shZbtb16), while transducing a non-targeting random shRNA as a negative control (shRandom). We then cultured transduced progenitors on OP9-DL1 monolayers in the presence of ILC- promoting cytokines, then analyzed them by flow cytometry after 10 days. Under these conditions, the majority of progenitors primarily activated *Bcl11b* and committed to the T cell lineage, for both shRandom and shZbtb16b-transduced cells, with a small percentage of cells entering the ILC2 or NK lineages (Lin^-^ICOS^+^DX5^-^Bcl11b^+^ or Lin^-^DX5^+^). Progenitors transduced with shZbtb16 constructs generated significantly more T cell precursors, as well as significantly fewer ILC2 and NK cells than their control counterparts transduced with shRandom constructs. These results demonstrate that PLZF is critical for promoting the divergence into the ILC lineage when *Bcl11b* activation is delayed and highlights the importance of the relative timing and order of transcription factor expression during multipotent progenitor cell fate decisions.

## DISCUSSION

Our results underscore the importance of developmental timing delays in controlling lineage decisions at developmental branch points. For ILC2 and T cell lineage specification from DN2 progenitors, we find that extended time delays in *Bcl11b* activation provide an opportunity for cells to divert from the canonical T cell lineage to an alternative ILC lineage. While ILCs are relatively rare in the adult thymus, recent evidence suggests that small numbers of resident ILC2s play a critical role in responding to thymic damage by promoting thymic epithelial differentiation to counteract thymic involution (Nevo et al., 2024). Thus, flexible and dynamic regulation of DN2- derived ILC2 populations balanced against T cell production in the thymus may be an important aspect of thymus functionality.

Our work suggests that delayed activation of *Bcl11b*, which is known to negatively regulate *Zbtb16* upon expression, may enable an extended period of *Zbtb16* expression prior to Bcl11b expression and result in a divergence of cells into the ILC pathway. Intriguingly, *Bcl11b* activation can still occur after up-regulation of the PLZF-dependent ILC program; in this case, there is a shift in transcriptional regulatory activity of Bcl11b from an pro-T cell program to a pro-ILC program (Hosokawa et al., 2018; Hosokawa et al., 2020). In an unperturbed system, the Bcl11b switch clearly occurs far more quickly on average than that of the PLZF-mediated switch and thus it is normally rare for Bcl11b-/PLZFhi ILC progenitors to emerge from DN2 progenitors (Figure 2D and Figure 5A). Our results demonstrate that the Bcl11b timing enhancer is responsible for upholding the rate of *Bcl11b* switching in relation to other PLZF-related switches operating in parallel and thus explains why removal of the timing enhancer increases ILC lineage generation at the expense of T cell output. This may also explain why others have found that under wild type conditions a small fraction of peripheral ILC2s appear to be derived from thymic DN3 progenitors but that fraction dramatically increases when the repressors of *Zbtb16*-E2A/HEB E proteins- are disrupted(Qian et al., 2019). Interestingly, *Zbtb16* may also be regulated by its own timing enhancer during natural killer T cell development, raising the possibility that competing timed switches controlling Bcl11b and PLZF activation may jointly regulate innate and adaptive fate output in the thymus.

The stochastic nature of the time delays in epigenetic switching may enable multipotent progenitors to generate multiple differentiated cell types in response to a common signal. While Notch signaling and its downstream regulators Gata3 and TCF-1 are essential for driving T cell lineage commitment, they also play critical roles in ILC development and also act upstream of the ILC regulators Id2 and PLZF. Thus, it is possible that these same instructive signals for T cell development also drive ILC development in the thymus. In this picture, the decision for whether to enter the T or the ILC pathway would be determined by the activation timing of *Bcl11b*, and possibly of PLZF, which may be also regulated by an enhancer-mediated stochastic epigenetic switch (Mao et al., 2017). As activation time delays would vary between cells due to the inherently stochastic nature of epigenetic switching, they could generate heterogeneity in fate outcomes to modulate relative cell population sizes, even amid uniform developmental signals.

While stochastic epigenetic timing control mechanisms are likely important for controlling differentiation outcomes in diverse contexts, they may be particularly useful for lineage control in the immune system, because immune cells are highly motile and thus less confined to specific niches with defined signaling environments compared to other cell types. We note that, despite its inherently stochastic nature, epigenetic switches can be highly controlled at multiple levels, including by TFs, *cis*-regulatory elements and chromatin modifying enzymes (Jacobsen et al., 2017; Kueh et al., 2016; Nguyen et al., 2021; Pease et al., 2021). These multiple layers of control would allow for precise tuning of differentiated cell numbers, particularly at the population level where variability in outcomes can be minimized due to averaging at large cell numbers (Nguyen et al., 2021).

In further studies, it would be important to more broadly investigate roles for epigenetic timing control in developmental gene regulation and decision making across diverse contexts. Timed epigenetic switches, when operating in the context of gene regulatory networks, have the ability to profoundly shape their dynamics and the developmental transitions they control. An understanding of these dynamics, both at an experimental and theoretical level, will help us more fully understand the developmental specification and evolutionary diversification of metazoan tissue size, shape and function.

## METHODS

### Animal models

C57BL/6 *Bcl11b^YFP/YFP^* mice (i.e. “WT” mice) and *Bcl11b^YFP^*^ΔEnh*/YFP*ΔEnh^ mice (i.e. “ΔTE”) were generated as previously described(Ng et al., 2018). *Bcl11b^YFP/YFP^* were crossed to *Bcl11b^YFP^*^ΔEnh*/YFP*ΔEnh^ and the resulting *Bcl11b^YFP/YFP^*^ΔEnh^ heterozygotes were used as breeding pairs to generate littermate match controls for primary cell analysis. All animals were bred and maintained at the University of Washington. All animal protocols were reviewed and approved by the Institute Animal Care and Use Committee at the University of Washington (Protocol No: 4397- 01).

### Cell line culture

Primary cells isolated from bone marrow were cultured on a OP9-DL1 monolayer stromal cells(Holmes and Zuniga-Pflucker, 2009) at 37℃ in 5% CO2 conditions with standard culture medium [80% MEM-alpha (Gibco), 20% Fetal Bovine Serum (Corning), Pen-Strep-Glutamine (Gibco)] supplemented with appropriate cytokines indicated below. Phoenix-Eco cells were cultured at 37℃ in 5% CO2 with standard culture medium [90% DMEM (Gibco), 10% Fetal Bovine Serum (Corning), Pen-Strep-Glutamine (Gibco)]. All cell lines were tested and found to be negative for mycoplasma contamination.

### Cell purification

Bone marrow progenitors used for *in vitro* T cell development assays were purified as previously described(Ng et al., 2018). Thymi and spleens from 3-week old mice were mechanically dissociated before re-suspending in Fc blocking solution with 2.4G2 hybridoma supernatant. Early stage thymocytes (ETP-DN4) and ILC populations were depleted of CD4 and CD8 thymocytes before analysis or sorting. Thymocyte suspensions were labeled with biotinylated CD4 and CD8 antibodies incubated with MACS Streptavidin Microbeads (Miltenyi, Biotec) in HBH buffer (HBSS (Gibco), 0.5% BSA (Sigma-Aldrich), 10 mM HEPES, (Gibco)), pre-filtered through cell separation magnet (BD Biosciences), and passed through a magnetic column (Miltenyi Biotec).

### *In vitro* differentiation of T cell progenitors

To generate double-negative (DN) T cells *in vitro*, thawed CD117-enriched bone marrow progenitors were cultured on OP9-DL1 stromal cell monolayers as described before(Ng et al., 2018) using standard culture medium [80% αMEM (Gibco), 20% Fetal Bovine Serum (Corning), Pen-Strep-Glutamine (Gibco)], grown at 37°C in 5% CO2 conditions]. All *in vitro* T cell generation cultures were supplemented with 5ng/mL Flt3-L and 5 ng/mL IL-7 (Peprotech). Experiments involving *in vitro* generation of NK and ILC populations (Figures 3, 4, and 5) were supplemented with 5ng/mL IL-7, 10ng/mL IL-2, 10ng/mL mSCF, 10ng/mL IL-15 (Peprotech).

### Flow cytometry and cell sorting

Fluorescence activated cell sorting (FACS) was used to isolate DN cells of interest with the following protocol. Bone marrow derived cell cultures were scraped and incubated in 2.4G2 Fc blocking solution and stained with anti-CD25 APC-eFluor 780 (Clone PC61.5, eBioscience) and with biotinylated antibodies against a panel of lineage markers (CD19, CD11b, CD11c, NK.1.1, Ter119, CD3ε, Gr-1 and B220 (BioLegend)). Stained cells were washed with HBH (Hank Balanced Salt Solution (HBSS) with 0.1% bovine serum albumin (BSA) and 10mM HEPES) and stained with streptavidin-PerCP/Cy5.5 (BioLegend). Stained cells were washed, resuspended in HBH, and filtered through a 40-um nylon mesh for sorting with a BD FACS Aria III (BD Biosciences) with assistance from the University of Washington Pathology Flow Cytometry Core Facility. A benchtop Attune NxT Flow Cytometer (ThermoFisher Scientific) was used to analyze primary and re-cultured thymocytes and acquired data were analyzed with FlowJo software (Tree Star).

### Single cell RNA-sequencing

Bone marrow progenitors attached to OP9-DL1 stromal cells were washed three times with PBS before scraping in HBH buffer and passing through a 70uM mesh filter to minimize OP9-DL1 stromal cell contamination. Cells were then washed with PBS and fixed in 4% paraformaldehyde for 15 minutes on ice. After fixation, cells were pelleted at 500xg for 3 minutes at 4°C before resuspending in 1mL of PBSR [PBS, pH 7.4, 1% BSA, 1% SUPERase-In RNase Inhibitor (ThermoFisher Scientific), and 1% 10mM DTT], pelleting again at 500xg for 5 minutes at 4°C and resuspending in PBSR. The Brotman Baty Institute (BBI) then performed the single cell transcriptome library construction following the sci-RNA-seq3 combinatorial indexing protocol as described in(Cao et al., 2019)

### Single cell RNA-sequencing analysis

Scanpy(Wolf et al., 2018) was used to first filter on cells with at least 200 genes and genes detected in at least 15 cells. Next, cells with greater than 2,000 gene expression counts or greater than 30% mitochondrial counts were removed. MIRA topic modeling was then applied with the most highly variable genes to describe each cell’s transcriptional state as a composition of co-regulated genes (i.e. topics) with variable weights(Lynch et al., 2022). The nearest neighbors distance matrix was computed with *n_neighbors* set to 40 and the Leiden algorithm was used to cluster cells with *resolution* set to 0.3. For visualization and down-stream analyses, cells were downsampled such that each genotype had equal numbers of cells in each starting state (12,353 for DN1 starting cells; 13,050 for DN2a starting cells; 5,631 for DN2b starting cells). Differential gene expression analysis between genotypes among the multipotent DN2 clusters (0, 1, 2, 4) was performed using the Wilcoxon rank sum test. Gene ontology analysis of the top 400 genes from the T cell and ILC topics (Table S1) was performed with the TopFun function of the TopGene suite(Chen et al., 2007).

### Retroviral vector generation and transduction

shRNA sequences targeting *Zbtb16* transcripts were joined to a U6 promoter and cloned into a Banshee-mCherry backbone as described previously(Kueh et al., 2016). The hairpin sequence is as follows: GGAAATGATGCAGGTAGATGA(anti-sense)-TTCG(loop)-TCATCTACCTGCATCATTTCC(sense). Retroviral particles were generated using the Phoenix-Eco packaging cell line. Viral supernatants were collected at 2 and 3 days after transfection and immediately frozen at -80°C. To infect bone marrow derived T cell progenitors, 33 μg/mL retronectin (Clontech) and 2.67 μg/mL of DL1-extracellular domain fused to human IgG1 Fc protein (a gift from I. Bernstein) were added in a volume of 250 μL per well in 24-well tissue culture plates (Costar, Corning) and incubated overnight. Viral supernatants were added the next day into coated wells and centrifuged at 2000 rcf for 2 hours at 32°C. Bone marrow derived derived T cell progenitors used for viral transduction were cultured for 6-7 days according to conditions described above, disaggregated, filtered through a 40-μm nylon mesh, and 10^6^ cells were transferred onto each retronectin/DL1-coated virus-bound well supplemented with 5 ng/mL mSCF (Peprotech), 5 ng/mL Flt3-L, and 5 ng/mL IL-7.

## ACKNOWLEDGEMENTS

The authors thank Xiaoping Wu, Ngoc-Han Nguyen, and Aurelio Silvestroni in the UW Pathology Flow Core for assistance with sorting. The authors also thank the Brotman Baty Institute (BBI) for the contribution to this work by way of performing single cell RNA- sequencing experiments and pre-processing the data. This work was funded by an NIH K99/R00 Pathway to Independence Award (NIH/ R00HL119638, to H.Y.K.), an NIH/NHLBI R01 (R01HL146478, to H.Y.K.), an National Science Foundation grant URoL EF-2021552 (to H.Y.K.); a John H. Tietze Foundation Stem Cell Scientist Award (to H.Y.K.), a Fellowship (to N.A.P.) from the University of Washington Institute of Stem Cell and Regenerative Medicine of the United States (to N.A.P.); and NIH F31 Fellowships F31 HL151090 (to N.A.P.)

## SUPPLEMENTARY FIGURE LEGENDS

**Fig. S1:**
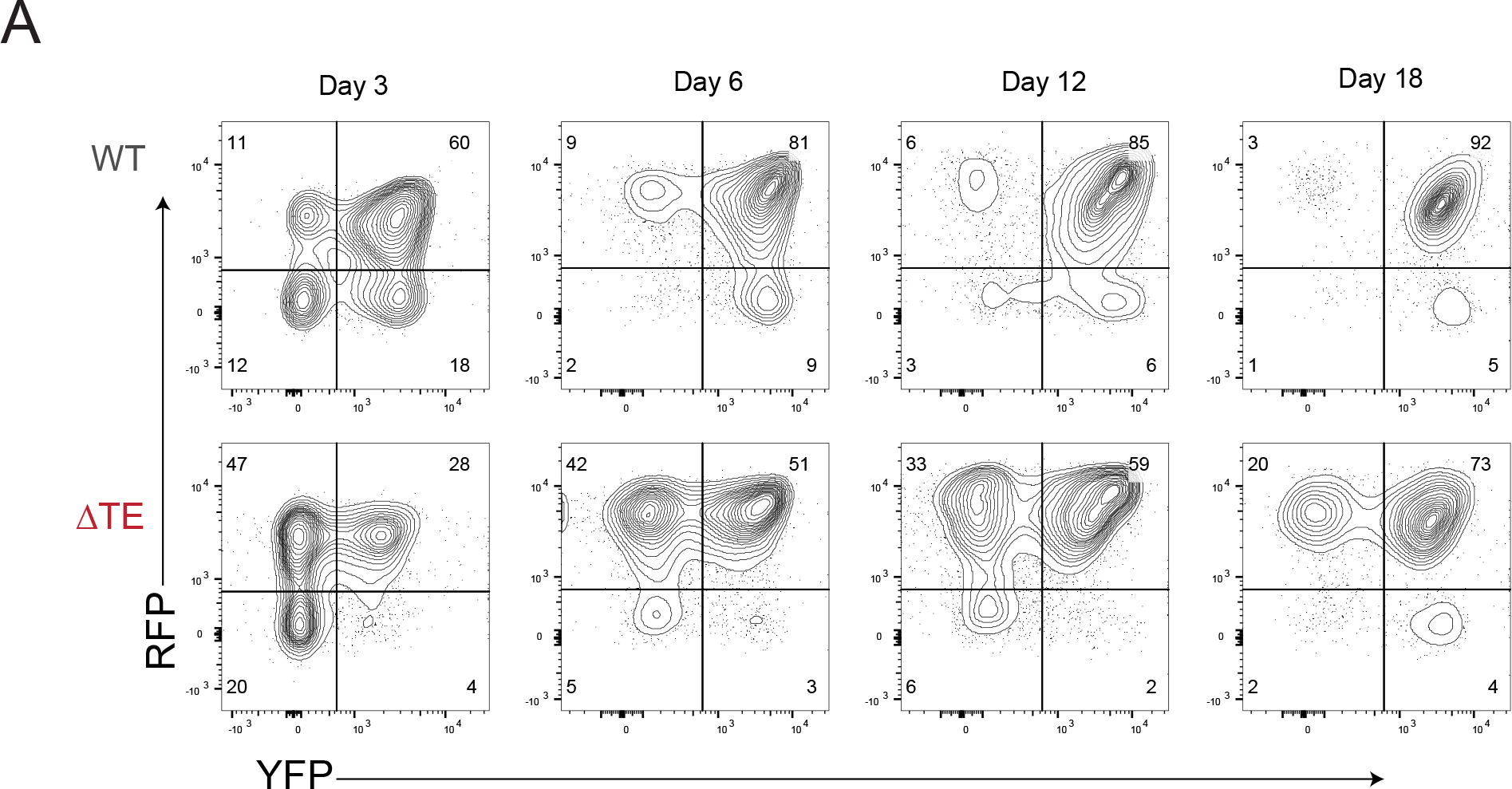
Timing enhancer regulates *Bcl11b* timing *in cis.* (A) Representative contour plots for Bcl11b-RFP and -YFP levels of purified DN2a progenitors re-cultured on OP9-DL1 cells.

**Fig. S2:**
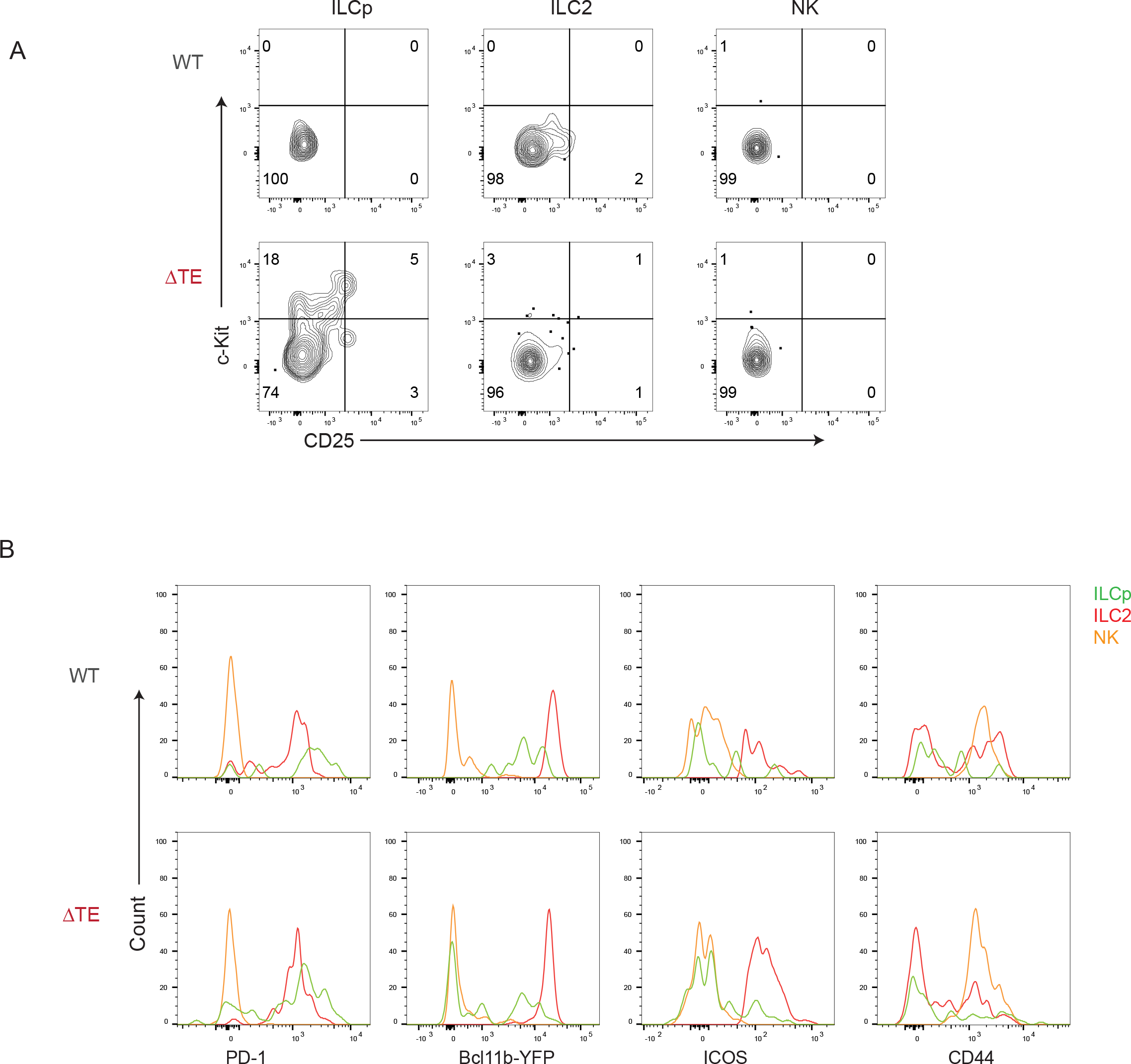
Immunophenotyping of thymic ILC populations. (A) Representative contour plots of thymic ILC subsets. (B) Representative histograms for ILC related markers.

**Fig. S3:**
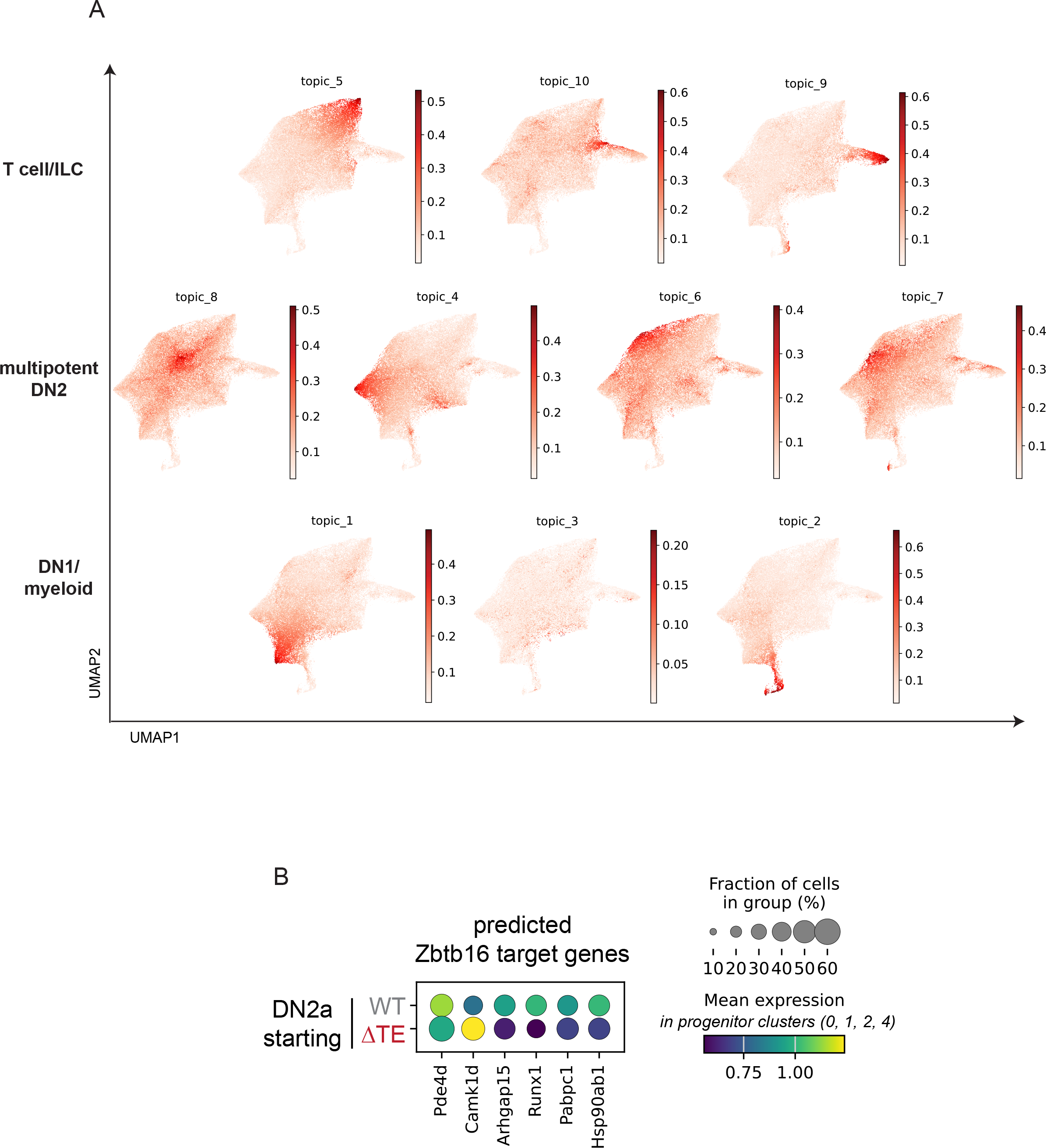
sci-RNA-seq gene topics and PLZF target gene expression. (A) UMAP representation of all cells colored by gene topic expression. (B) Normalized expression levels of selected top CellOracle predicted PLZF target genes across in DN2a-starting cells in the progenitor clusters.

## Notes

### Competing Interest Statement

The authors have declared no competing interest.

